# Theoretical foundation of the RelTime method for estimating divergence times from variable evolutionary rates

**DOI:** 10.1101/180182

**Authors:** Koichiro Tamura, Qiqing Tao, Sudhir Kumar

## Abstract

RelTime estimates divergence times by relaxing the assumption of a strict molecular clock in a phylogeny. It showed excellent performance in estimating divergence times for both simulated and empirical molecular sequence datasets in which evolutionary rates varied extensively throughout the tree. RelTime is computationally efficient and scales well with increasing size of datasets. Until now, however, RelTime has not had a formal mathematical foundation. Here, we show that the basis of the RelTime approach is a relative rate framework (RRF) that combines comparisons of evolutionary rates in sister lineages with the principle of minimum rate change between an evolutionary lineage and its descendants. We present analytical solutions for estimating relative lineage rates and divergence times under RRF. We also discuss the relationship of RRF with other approaches, including the Bayesian framework. We conclude that RelTime will be also useful for phylogenies with branch lengths derived not only from molecular data, but also morphological and biochemical traits.

## INTRODUCTION

The inference of divergence times is usually accomplished by assuming a constant rate throughout the tree (a strict molecular clock) or by relaxing the strict molecular clock (Ho and Duchêne 2014; Kumar and Hedges 2016; dos Reis et al. 2016). Bayesian approaches are widely applied for relaxed clock analyses. They require specification of a probability distribution of evolutionary rates in the tree (e.g., lognormal distribution), as well as presence or absence of rate autocorrelation among branches. In contrast, the RelTime approach does not require specification of such priors, and produces relative node ages, which can then be transformed into absolute dates by using calibration constraints for one or more nodes (Tamura et al. 2012; Tamura et al. 2013). RelTime has been found to perform well for estimating divergence times in analyses of many large empirical datasets (Mello et al. 2017) and simulated datasets (Tamura et al. 2012; Filipski et al. 2014).

RelTime’s computational speed (**Fig. 1a**) and accuracy has led to its use for estimating node divergence time for large datasets (Tamura et al. 2012; Mahler et al. 2013; Bond et al. 2014; Bonaldo et al. 2016). However, a mathematical foundation for the RelTime method has not yet been provided. A theoretical foundation is needed not only to understand basic properties and assumptions of RelTime, but also to reveal its relationship with other molecular dating methods (Ho and Duchêne 2014; Kumar and Hedges 2016; dos Reis et al. 2016). In the following, we present the theoretical foundation of the RelTime method. We also assess the absolute performance of RelTime and compare its performance with other methods by analyzing data generated using computer simulations in which sequences were evolved according to three different branch rate models: independent (Drummond et al. 2006), autocorrelated (Kishino et al. 2001), and hybrid (Beaulieu et al. 2015).

**Figure 1.**
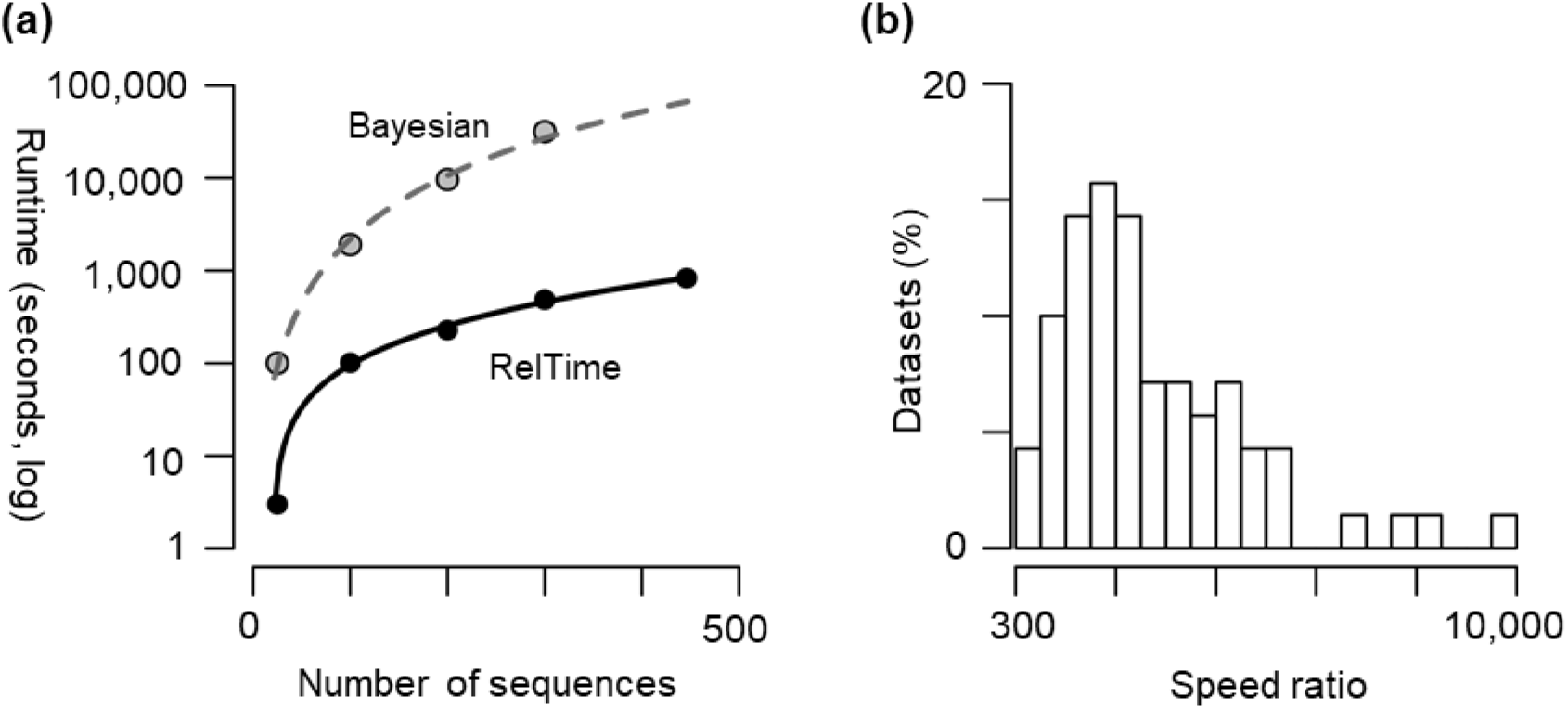
(**a**) Computational time taken by RelTime and MCMCTree (Bayesian method) to estimate divergence times for datasets containing increasing number of sequences (*n*). The tested sequence alignment consisted of 4,493 sites in which sequence evolution was simulated with extensive rate variation (RR50 data from Tamura *et al*. (2012)). RelTime’s speed advantage increases with data volume by *O*(*n*_2_). (**b**) Distribution of computation speed ratio of RelTime to MCMCTree for 70 datasets, each containing 100 ingroup sequences that were simulated as described in the **Materials and Methods**.

## MATHEMATICAL THEORY

### Theoretical analysis for a phylogeny with three taxa and an outgroup

We begin with the simplest case in which the phylogeny contains a clade with three ingroup taxa (subtree at node 5) and one outgroup taxon (**Fig. 2a**). In this tree, *b*_1_ and *b*_2_ represent the amount of evolutionary change that has occurred in lineages emanating from node 4 and leading to taxon 1 and taxon 2, respectively. We assume that taxon 1 and 2 are sampled at the same evolutionary time (*t*_1_ = 0 and *t*_2_ = 0), which is usually the case in phylogenetic analysis of data sampled from living species. The contemporaneous sampling of data produces sampling times equal to 0, which serve as calibration points (Tamura et al. 2012). By using branch lengths in this phylogeny (*b*’s), we estimate the relative evolutionary rates (*r*’s) for all the lineages as well as relative divergence times (*t*’s). Here, a lineage refers to a branch and all the taxa (and branches) in the descendant subtree, e.g., lineage *a* in **Fig. 2a** contains two taxa (1 and 2) and three branches with lengths *b*_1_, *b*_2_, and *b*_4_.

**Figure 2.**
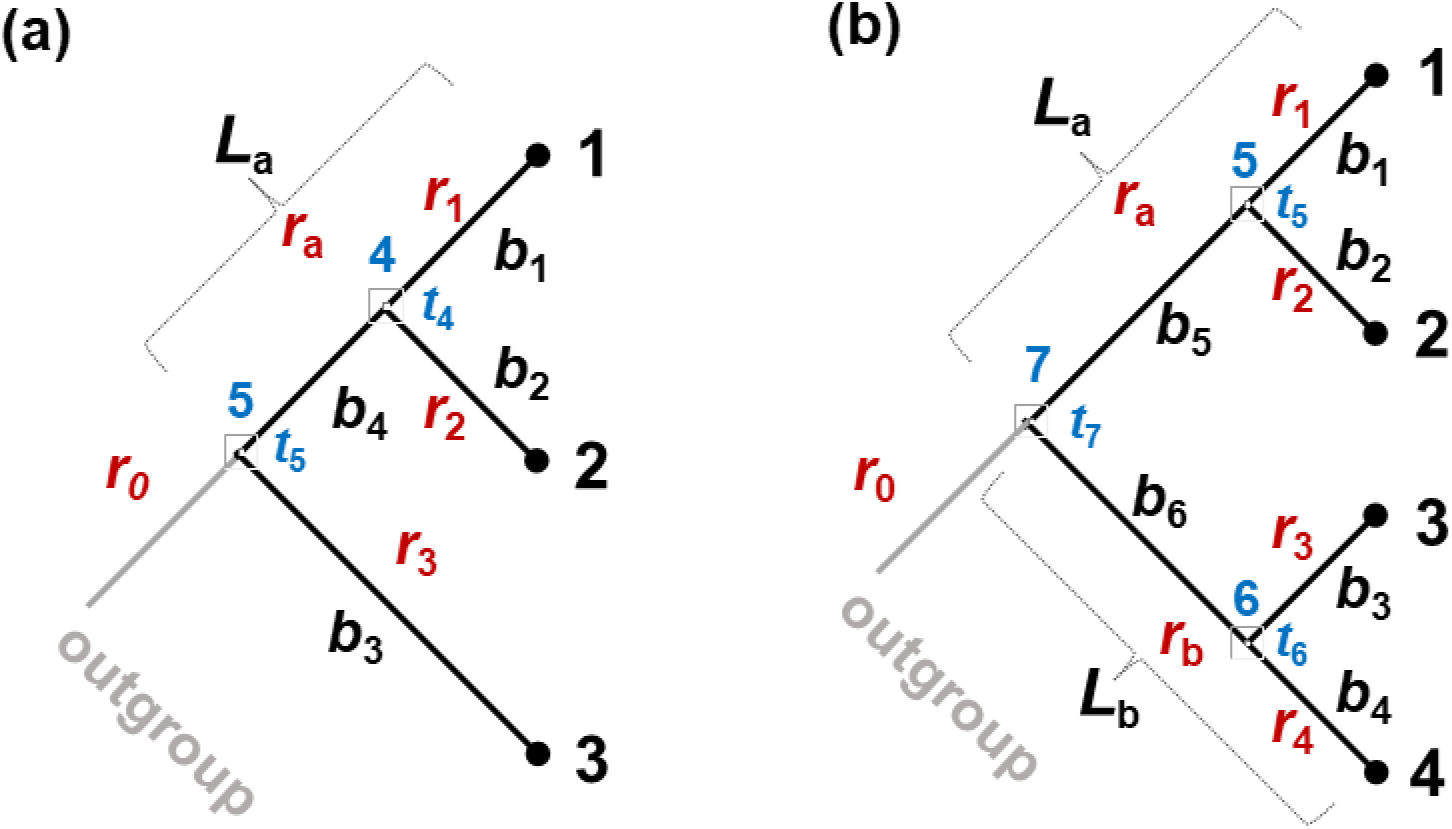
The relative rate framework for RelTime method. (**a**) A tree containing three ingroup sequences with an outgroup. Branch lengths are *b*_i_’s and lineage rates are *r*_i_’s. *L*_a_ = *b*_4_ + ½(*b*_1_ + *b*_2_). (**b**) The case of four ingroup sequences with an outgroup. Here, *L*_a_ = *b*_5_ + ½(*b*_1_ + *b*_2_). *L*_b_ = *b*_6_+ ½(*b*_3_ + *b*_4_), when using the arithmetic mean. See Figure 3 and its legend for a simple procedure outlining the calculation of relative rates using RRF.

The following system of equations formalizes the RelTime approach mathematically by linking relative rates for lineages (*r*_i_) with branch lengths (*b*_i_) in **Fig. 2a**. Here,

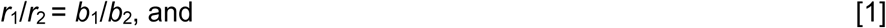

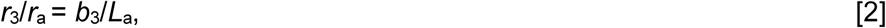
where, *L*_a_ is depth of node 5 on lineage *a*, which contains taxon 1 and 2; *L*_a_ = *b*_4_ + ½(*b*_1_ + *b*_2_). We set

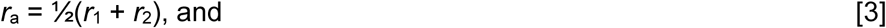

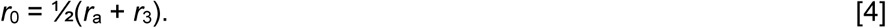
The setting of equalities in [3] and [4] leads to preference for the minimum rate change between the stem branch originating at node 5 and the descendant subtree originating at node 4. The selection of lineage rate *r*_a_ will be constrained by equations [3] and [4], which relaxes the strict molecular clock.

Because all the rates are relative, one unknown is reduced by setting the group rate at the most recent common ancestor of the ingroup node to be 1, *i.e.*,

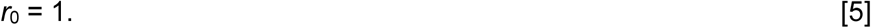
We solve for each *r*_i_ by using equations [1] – [5] and get

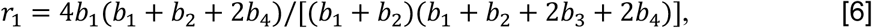

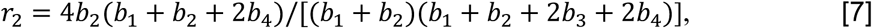

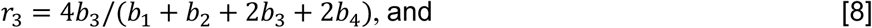

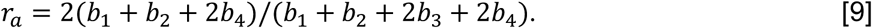
The estimates of relative lineage rates and lengths produce an ultrametric tree with relative times for node 4 (*t*_4_) and node 5 (*t*_5_):

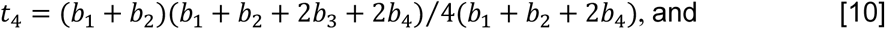

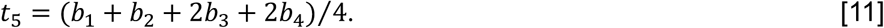
The above equations ([1] – [11]) constitute the relative rate framework (RRF) for phylogenies with three taxa and one outgroup. These equations yield point estimates for lineage rates and node ages. Because branch lengths have variances, the resulting lineage rates estimates have variances. The variance of relative node ages will be a function of branch length variances and the variance contributed by evolutionary rate differences among branches (see **Discussion**).

### Theoretical analysis for phylogenies containing four taxa and an outgroup

We need to estimate six evolutionary rates (*r*_1_ – *r*_4_, *r*_a_, and *r*_b_) using six branch length estimates (*b*_1_ – *b*_6_) for a phylogeny containing four ingroup taxa (**Fig. 2b**). Following the above approach, we write a set of seven equations:

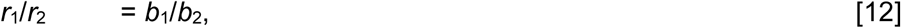

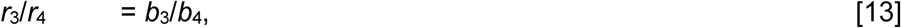

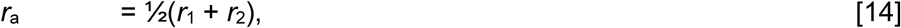

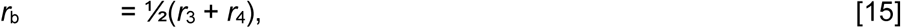

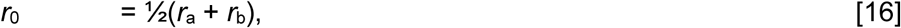

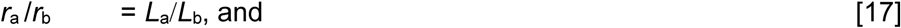

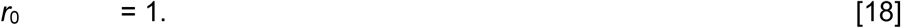
Here, *L*_a_ = *b*_5_ + ½(*b*_1_ + *b*_2_) and *L*_b_ = *b*_6_ + ½(*b*_3_ + *b*_4_). Equations [14] – [16] above lead to a preference for minimum changes in rates between ancestor-descendant lineage pairs. In this system, *r*_i_ ‘s are not required to be equal to one another and the rate assignments are constrained to respect the rate ratios among lineages (equations [12], [13], and [17]), which relaxes the molecular clock.

We solve for each *r*_i_ from equations [12] – [18] to estimate relative rates and node ages.

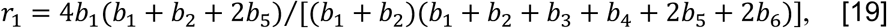

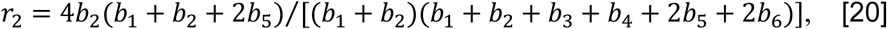

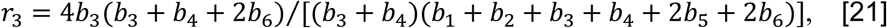

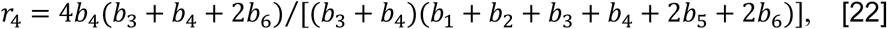

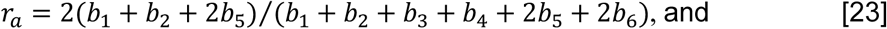

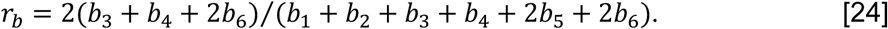
The estimates of relative node ages *t*_5_, *t*_6_, *t*_7_ for nodes 5, 6, and 7, respectively, are:

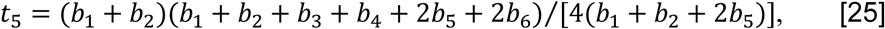

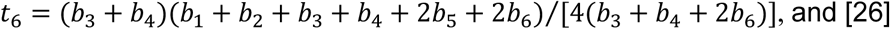

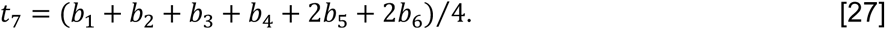
The above equations ([19] - [27]) establish RRF for the RelTime approach for a tree containing four taxa and one outgroup. As previously mentioned, point estimates of node ages and lineage rates have variances because branch lengths have variances and evolutionary rates are not equal among lineages (see **Discussion**).

### Relative rate framework with geometric means

In both the original RelTime approach (Tamura et al. 2012) and the mathematical formulations above, we considered an arithmetic mean when averaging branch lengths to minimize evolutionary rate changes. This approach does not assume an equal rate, but is rather a natural way to calculate node depths by averaging branch lengths. We have now developed analytical formulas for an alternative RRF in which the geometric mean is used, which balances the rate changes between two descendant lineages. For example, if *b*_1_ = 1 and *b*_2_ = 4 in **Fig. 2a**, then the arithmetic mean will give 2.5. Thus, evolutionary rate *r*_1_ is 2.5 times slower and *r*^2^ is 1.6 times faster when compared to rate of on the ancestral branch (*b*_4_). The difference in rate change (between 2.5 and 1.6, in the present case) becomes larger as the difference between *b*_1_ and *b*_2_ becomes larger when using the arithmetic mean. In contrast, the geometric mean would give 2.0, which results in two times slower rate in *b*_1_ and two times faster rate in *b*_2_, as compared to the ancestral branch. That is, the difference in rate between the ancestral and descendant lineages is always equal for sister lineages when using the geometric mean, which is not the case if the arithmetic mean is used.

Using a geometric means approach, we obtain the following analytical formulas for a phylogeny containing three ingroup taxa (**Fig. 2a**):

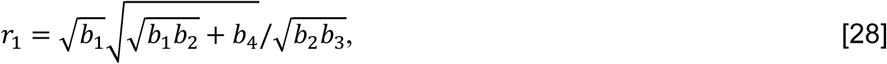

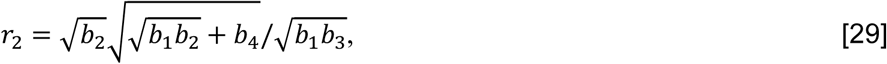

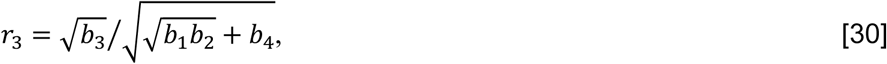

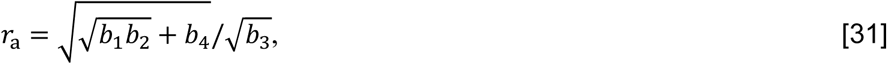

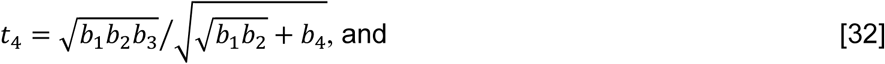

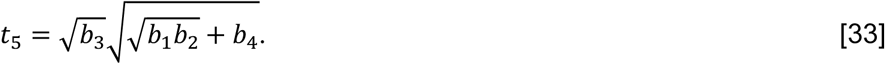
For the 4-taxon case in **Fig. 2b**, the equations are as follows:

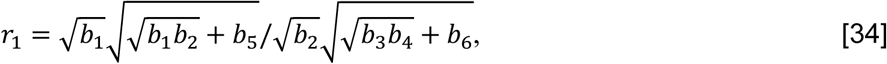

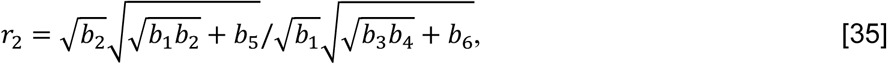

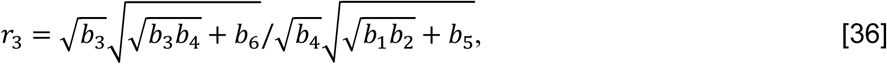

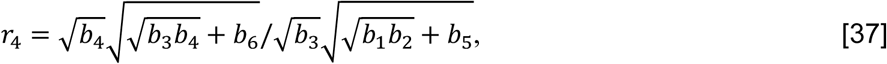

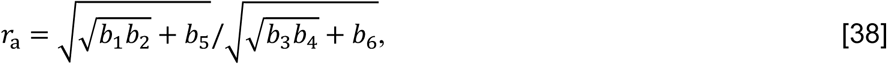

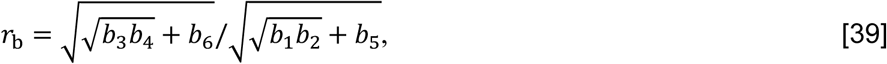

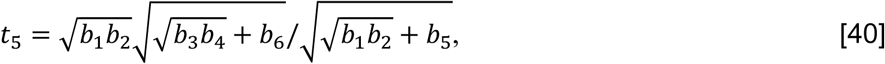

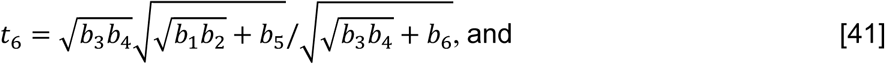

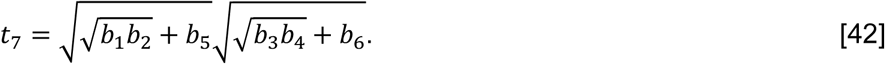

### Relative rate framework for a general case

Now we consider a general case of a phylogeny with more than four ingroup taxa. In this case, RelTime applies RRF in a bottom-up approach, starting from the tips (external branches) of the phylogeny and moving towards the root. In the first step, we generate local relative lineage rates for the subtrees containing three and four taxa (subtree *x* and *y*, respectively) in a phylogeny containing eight taxa and an outgroup (**Fig. 3**). Subtree *y* contains four taxa, so we apply equations [34] – [39] to generate relative rates. Equations [28] – [31] will be used to estimate rates for clade *x* which contains three ingroup taxa.

**Figure 3.**
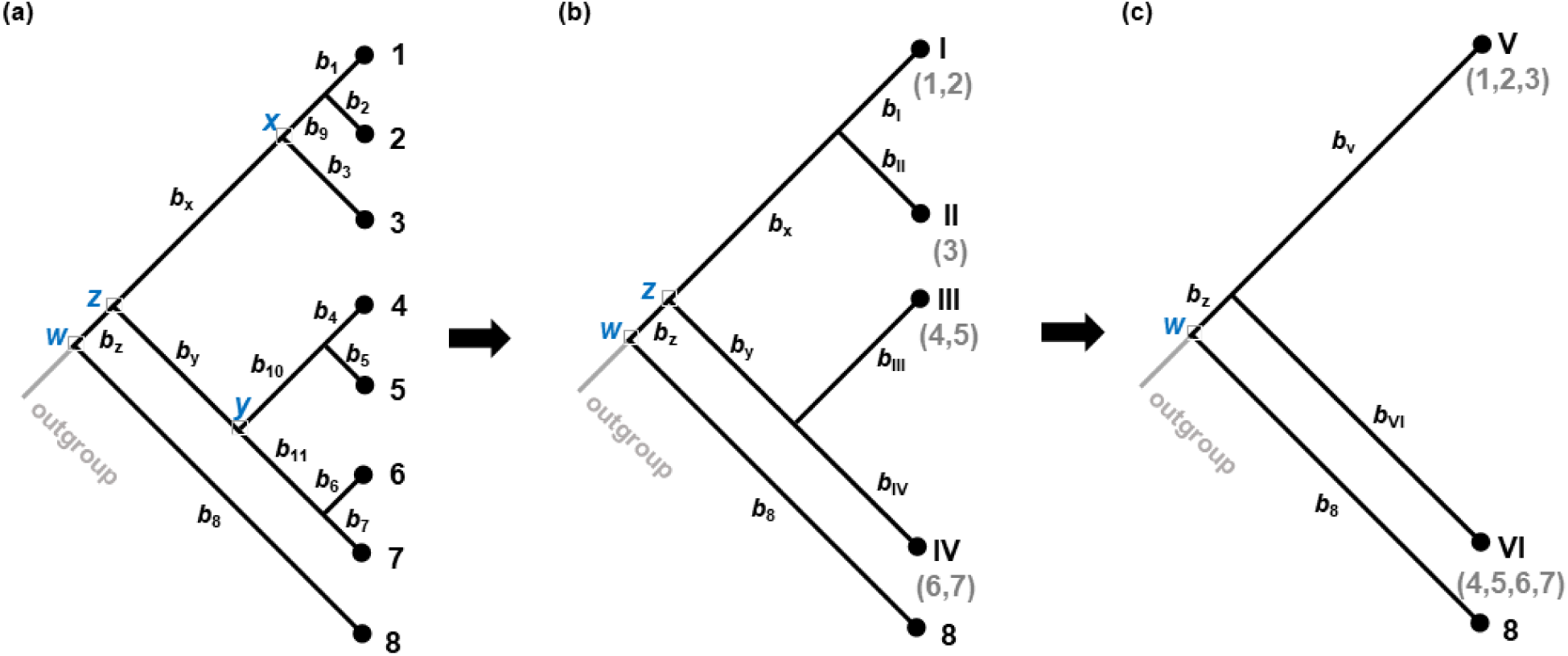
Calculating relative rates in a phylogeny. (**a**) A phylogeny containing eight ingroup sequences (taxa) that consists of three taxa nodes (*x* and *w*) and four taxa nodes (*y* and *z*). (**b**) Reduced phylogeny and branch lengths after applying RRF to nodes *x* and *y*. (**c**) Final phylogeny after applying RRF to node *z* in panel **b**, which produces node *w* in a three taxa configuration. After applying RRF to node *w*, a pre-order traversal scales descendant lineage rates by multiplying them by their ancestral lineage rates to generate final relative rates. The ingroup root node (*w*) has an average relative lineage rate equal to 1. Multi-sequence taxa are indicated by Roman numerals.

In the next step, we consider the parent clade (*z*) that has four taxa: composite taxon I consisting of two taxa (1 and 2), taxon II consisting of one taxon (3), composite taxon III consisting of two taxa (4 and 5), and composite taxon IV containing two taxa (6 and 7) (**Fig. 3b**). We estimate branch lengths (*b*_I_, *b*_II_, *b*_III_, and *b*_IV_) by using the geometric means: 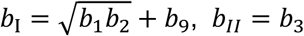, 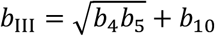, and 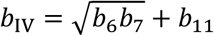. Then, we use equations [34] – [39] to compute relative rates for all the lineages using these branch lengths. In bigger phylogenies, this process is carried out for every parent node in a post-order traversal. In the current example, node *w* is the common ancestor of all the ingroup taxa, which is in a three taxa configuration (**Fig. 3c**). We now have 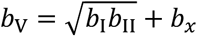 and 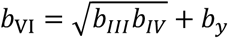 and apply equations [28] – [31]. At this stage, we have local (relative) lineage rate estimates for nodes in the ingroup tree. Finally, all the rates in the tree are computed by multiplying descendant lineage rates by their respective ancestral lineage rates in order to generate final relative rates such that the ingroup root node (e.g., node *w* in **Fig. 3c**) has an average relative lineage rate equal to 1.

## RESULTS

We evaluated the performance of RRF for correctly estimating lineage rates and divergence times by analyzing data generated using computer simulation in which sequences were evolved according to the autocorrelated rate [AR] model (Kishino et al. 2001), the independent rate [IR] model (Drummond et al. 2006), as well as a model that contains multiple distributions of rates (hybrid rates [HR]). We present results from the analysis of a collection of small datasets (three ingroup sequences; AR and IR models) and two collections of large datasets: one containing 100 ingroup sequences (AR and IR models) and another containing 91-ingroup sequences (HR model).

### Analysis of small datasets (phylogeny with three ingroup sequences)

We first tested the accuracy of divergence times estimated via RRF by analyzing datasets containing three ingroup sequences that were generated through computer simulations. In these datasets, sequences evolved according to either an autocorrelated rate (AR) model or an independent rate (IR) model (see **Materials and Methods** for details). We used RRF with geometric means and compared the modeled (true) lineage rates, estimated lineage rates, as well as node ages (**Fig. 4**). The lineage rates produced by RRF were similar to the true rates for datasets that evolved under the AR model (**Fig. 4a**); the relationship showed a linear regression slope of 1.0. RRF estimates for external lineages, which consist of three sequence each, were similarly good (blue circles, **Fig. 4a**). The relationship was also strong for internal branches, but with greater dispersion; *r*^2^ = 0.29 for red circles as compared to 0.68 for blue circles in **Fig. 4a**. RRF performance for AR datasets was similar to that observed for IR datasets (**Fig. 4b**). Overall, the success in estimating lineage-specific evolutionary rates translates into robust estimates of relative times for AR (**Fig. 4c**) and IR datasets (**Fig. 4d**).

**Figure 4.**
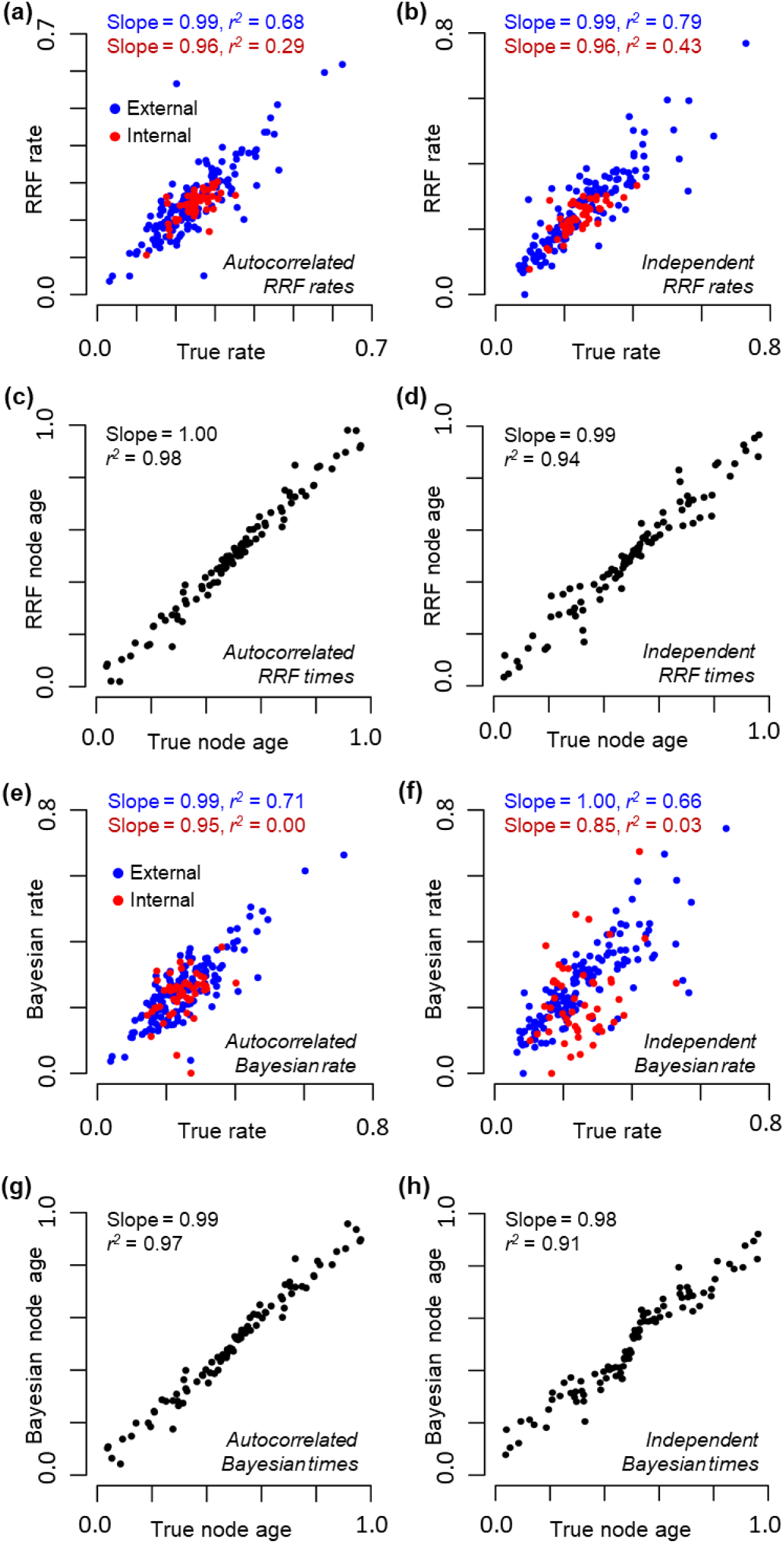
Performance of RRF and MCMCTree Bayesian analyses for three ingroup sequences with an outgroup (topology in Fig. 2a). RRF lineage rate estimates are compared with the true lineage rate estimates for sequences evolved under (**a**) autocorrelated and (**b**) independent rate models. Blue circles represent external lineages (single taxon, *r*_1_, *r*^2^, and *r*_3_) and red circles represent the internal lineage (*r*_a_). RRF estimates of divergence times for (**c**) autocorrelated rate and (**d**) independent rate datasets. Bayesian (MCMCTree) estimates of branch rates are compared with the true branch rates for sequences evolved under (**e**) autocorrelated and (**f**) independent rate models. Blue circles indicate external branches (*r*_1_, *r*^2^, and *r*_3_) and red circles show the internal branch (*r*_4_). Bayesian estimates of divergence times for (**g**) autocorrelated rate and (**h**) independent rate datasets. Each panel contains results from 50 simulated datasets. All rates and divergence time estimates are normalized to allow direct comparison between true and estimated values. Regression slope and correlation coefficient (*r*^2^) are shown for each panel.

Bayesian methods produce branch-specific rates under a given statistical distribution of rates (e.g., lognormal), which differs from RRF where relative lineage rates are considered. Therefore, we compared branch rates derived using Bayesian methods with their true values to evaluate the Bayesian based analyses. We provided correct priors (based on simulation parameters) and conducted the analyses using MCMCTree software (Yang 2007). Bayesian branch rate estimates showed a more diffuse relationship with true rates in internal branches for AR datasets (**Fig. 4e**; *r*^2^ = 0.00) as compared with RRF (**Fig. 4a**; *r*^2^ = 0.29). Fundamental reasons underlying these patterns are presented in the **Discussion** section. These patterns notwithstanding, Bayesian estimates of times for AR datasets showed a slope of 1.0 with true time estimates (**Fig. 4g**), which means that node age estimates are generally robust to difficulties in estimating branch-specific rates. This robustness was also seen for IR datasets, where Bayesian branch rates showed a more diffuse relationship with the true rates (**Fig. 4f**), but estimated times showed a slope close to 1 with a high *r*^2^ (0.91). Overall, both Bayesian and RRF approaches showed similar or lower accuracy in estimation of rates and node ages for IR datasets as compared to AR datasets, potentially because rate independence requires the estimation of a greater number of free parameters.

### Analysis of large datasets (phylogeny with 100 ingroup sequences)

We next analyzed datasets consisting of 100 ingroup sequences, which were evolved over a range of empirical rate variation parameters. As observed for datasets containing only three ingroup sequences, RRF lineage rate estimates were highly correlated with the true rates for AR datasets (**Fig. 5a**). Lineage rate correlations were generally lower for IR datasets and these correlations were higher for external (tip) lineages (**Fig. 5b**). Importantly, distributions of RelTime node age estimates were centered close to 1 for both AR and IR datasets (**Fig. 5c**).

**Figure 5.**
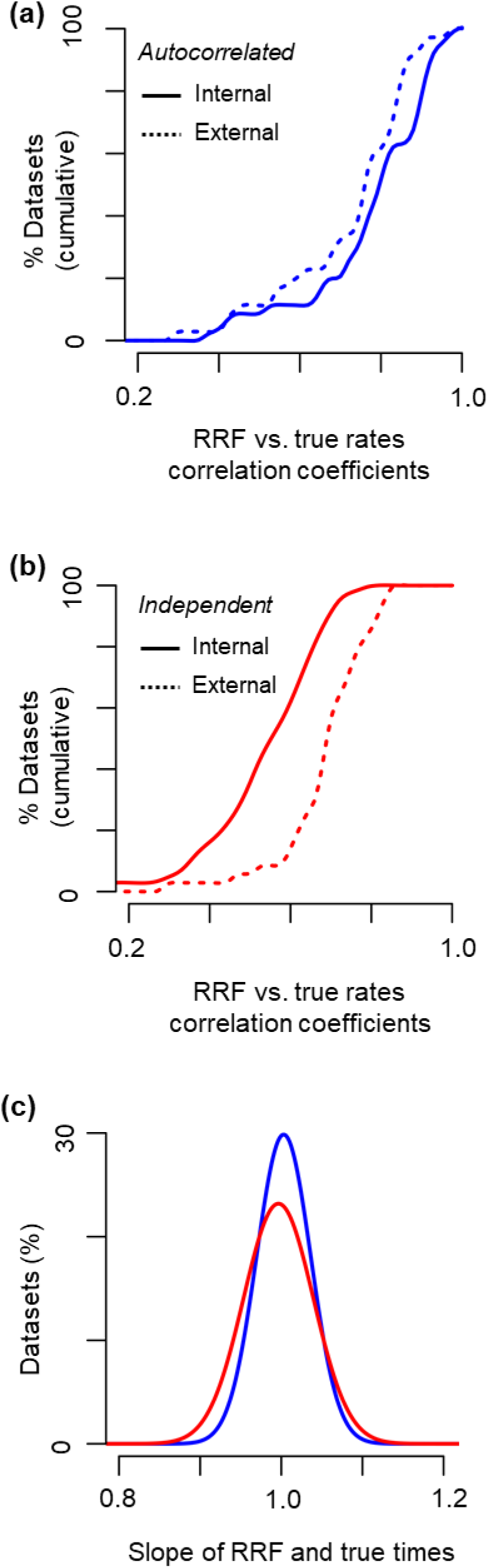
Performance of RRF in the analysis of datasets with 100 ingroup sequences and an outgroup. Fraction of datasets for which RRF inferred lineage rates are correlated with true rates at different levels of correlation for datasets simulated with (**a**) autocorrelated rates (**b**) independent rates. Dotted lines represent external branches and solid lines indicate the internal branches. (**c**) Distribution of the linear regression slopes of RRF estimates and true times for different datasets. The results presented are from analyses of 35 datasets in which sequences evolved with autocorrelated branch rates (blue lines) and another 35 datasets that were evolved with independent branch rates (red lines). In regression analysis, the intercept was set to zero, because the estimated node age is expected to be zero when the true node age is zero. The regression slopes generated with and without this assumption produced similar patterns.

### Analysis of hybrid rates datasets (phylogeny containing 91 ingroup sequences)

We also examined the performance of RRF in an analysis of simulated data from Beaulieu et al. (2015), who simulated two lognormal distributions (hybrid rate model) for an angiosperm phylogeny in which herbaceous clades exhibited higher and more variable evolutionary rates than woody clades (**Fig. 6a**). They reported that single-model Bayesian methods produced considerably more ancient date estimates for the divergence of herbaceous and woody clades.

**Figure 6.**
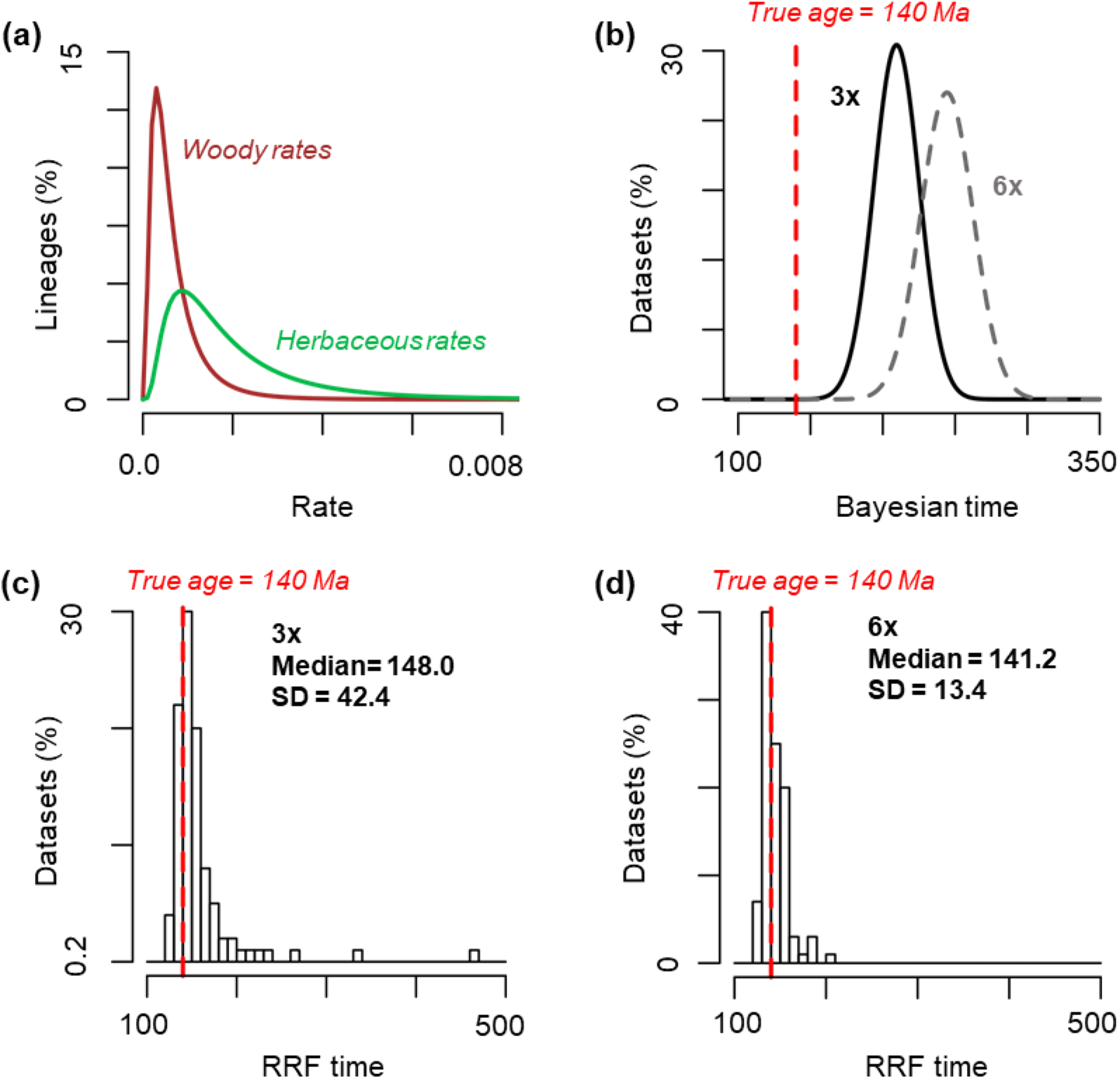
(**a**) Hybrid distribution of rates for branches leading to woody taxa (brown) and herbaceous taxa (green), with the former evolving three times (3x) slower than the latter. (**b**) Bayesian estimates reported by Beaulieu et al. (2015) when the rate difference between clades was three-fold (3x, solid line) and six-fold (6x, dashed line), with the simulated angiosperm age of 140 million years ago shown by a red line. RelTime estimates of angiosperm age for Beaulieu et al. (2015)’s alignments with (**c**) 3x mean rate difference and (**d**) 6x mean rate difference. Median and standard deviation for age estimates are shown. Beaulieu et al. (2015) simulated 100 replicates (1,000 bases) under a GTR model for each scenario. Bayesian analyses were conducted using a single uncorrelated lognormal rate prior in Beaulieu et al. (2015). The same alignments, topology and ingroup calibrations were used in RRF analyses.

This overestimation of divergence time became more severe as the difference between the two rates increased (**Fig. 6b**). Application of RRF produced divergence time estimates that were much closer to true times (**Fig. 6c** and **6d**), which shows that RRF can be useful in cases where the rate distribution differs among clades (Smith and Donoghue 2008; Dornburg et al. 2011; Beaulieu et al. 2015) or when clocks are local (Drummond and Suchard 2010; Crisp et al. 2014).

As a further example, Tamura et al. (2012) found that RelTime produced accurate time estimates in simulations with a very large number of sequences even when one clade possessed accelerated evolutionary rates, where penalized likelihood methods did not perform as well. In general, we expect that the limitations of single-model Bayesian analyses will be overcome by local clock methods (Drummond and Suchard 2010; Hӧhna et al. 2016; Lartillot et al. 2016), but the computational time required to analyze even modestly sized datasets via these approaches can be prohibitive. So, the current RRF approach, which does not assume a specific model for rate variation, may be suitable for such data in its current implementation or as a foundation for future methodological refinement.

## Discussion

We have presented a mathematical foundation for the relative rate framework (RRF) underlying the RelTime method. In the following, we compare RRF with other approaches for estimating divergence times and present the motivation behind RRF.

### Lineage rates versus branch rates

In RRF, evolutionary rate heterogeneity in a phylogeny is considered by comparing rates between sister lineages emanating from internal nodes in a phylogeny. A lineage rate is an average of all the lineage rates that belong to that lineage, including all the descendants (e.g., node *x* in **Fig. 3**). This focus on comparing lineage rates is fundamentally different from the comparison and modelling of branch rates in other approaches. For example, the penalized likelihood methods consider differences in rates between ancestral and descendant branches (Sanderson 1997; Sanderson 2003), and Bayesian methods model branch rates to share a probabilistic distribution (e.g., lognormal distribution).

The distinction between lineage and branch rates complicates direct comparisons of evolutionary rates produced by using RelTime and Bayesian methods, except for external (tip) branches for which the lineages consist of only one branch. In this case, RelTime and Bayesian estimates of rates show similar trends (**Fig. 4**, blue circles). This trend was also observed in the analysis of 100 sequence datasets, where the correlation of the estimated external branch rates with true branch rates was high for both RelTime (median *R*^2^ = 0.76 and 0.69) and Bayesian (*R*^2^= 0.72 and 0.86) analyses of AR and IR datasets, respectively.

For internal branches, computer simulations showed greater similarity between RRF estimates of lineage rates and the true lineage rates (**Fig. 4a** and **4b**) as compared to the similarity observed between Bayesian estimates of branch rates and the true branch rates (**Fig. 4e** and **4f, respectively**). This was also the case in the analysis of 100 sequence datasets: internal branch rates from Bayesian methods were less strongly correlated with the true rates (0.42 and 0.36 for AR and IR datasets, respectively), as compared to those observed for lineage rates from RRF (0.79 and 0.55 for AR and IR datasets, respectively).

These trends are due to the fact that the estimate of a branch rate is a function of two time estimates: one for the ancestral node and another for the descendant node. For example, the variance of the rate on branch with length *b*_4_ in **Fig. 2a** is a function of the variance of two time estimates (*t*_4_ and *t*_5_), in addition to the variance of *b*_4_. In contrast, the variance of a lineage rate (e.g., *r*_a_ in **Fig. 2a**) is a function of the variance of only one time estimate (*t*_5_), in addition to the lineage depth (*L*_a_), because the other time point is zero in contemporary sequence sampling. Thus, branch rates are estimated with greater variance than lineage rates, which results in lower correlations seen for Bayesian approaches.

### Underlying evolutionary rate model in RRF

RRF exploits the fact that the estimation of the ratio of lineage rates at any node in a phylogeny is independent of that node’s age. For example, the ratio of evolutionary rates at node 4, *r*_1_/*r*^2^, does not depend on *t*_4_, and it is clear that the rate of evolution is higher in the lineage leading from node 4 to taxon 2 than to taxon 1 (*r*^2^ > *r*_1_), because *b*_2_ is longer than *b*_1_ in **Fig. 7a**. In fact, we can estimate *r*_1_/*r*^2^ (= *b*_1_/*b*_2_) without knowing anything about the probability distribution of evolutionary rates throughout the tree. Similarly, the other rate ratio in this tree does not depend on knowledge of distribution of rates among branches, it is simply [(*b*_1_ + *b*_2_)/2 + *b*_4_]/*b*_3_ when using the arithmetic mean and 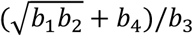 when using the geometric means.

**Figure 7.**
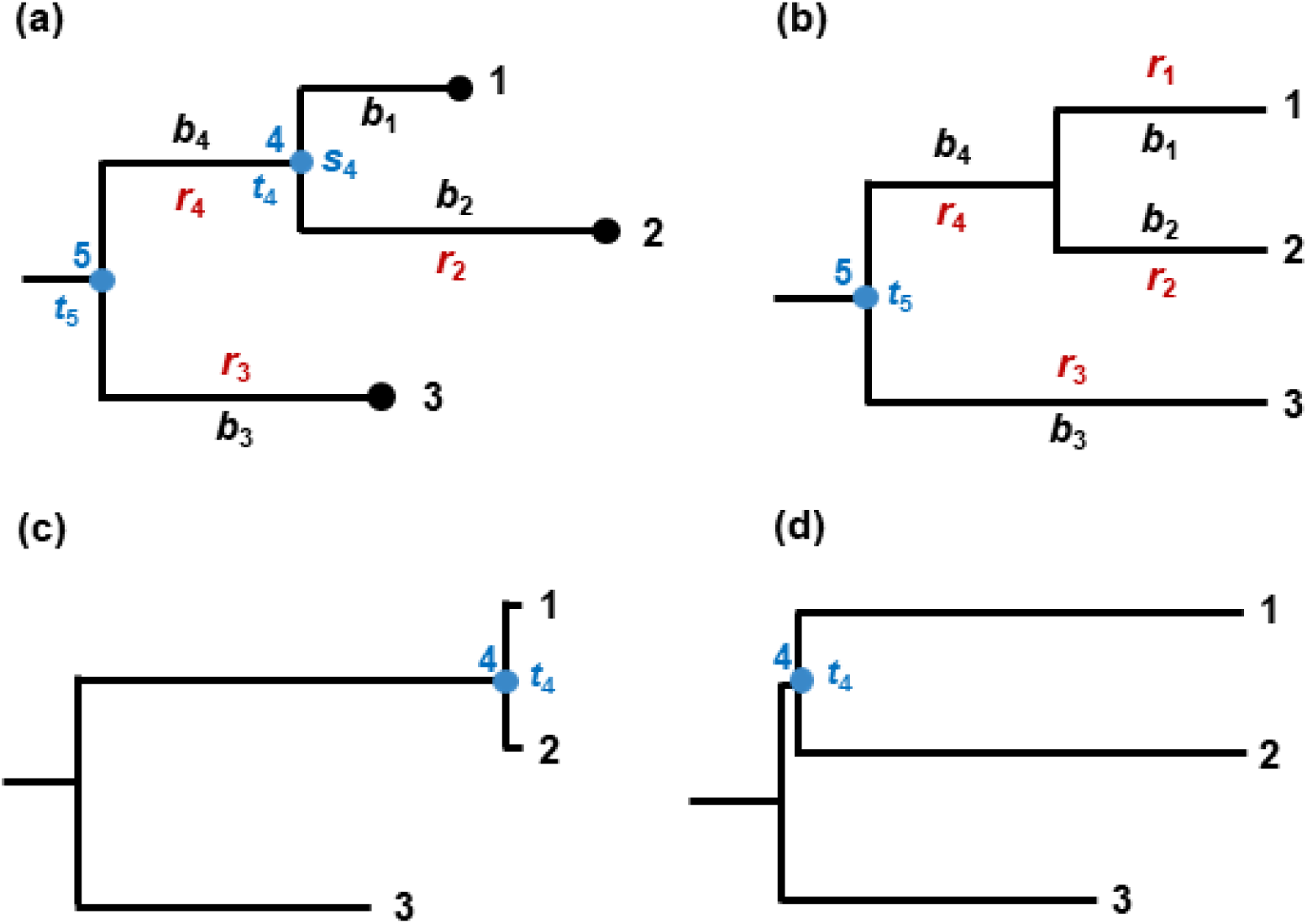
A phylogenetic tree of three taxa (1, 2 and 3). (**a**) Original phylogenetic tree with the observed branch lengths (*b*’s), which are necessary to estimate node times (*t’*s) shown in panel (**b)**. Evolutionary trees where the rate for the subtree containing taxon 1 and 2 (*s*_4_) is much (**c**) faster or (**d**) slower than that of its ancestral branch (*r*_4_).

However, the node-by-node specification of relative lineage rates is not sufficient to estimate relative times *t*_4_ and *t*_5_. For that, we need to know the relationship of subtree rate *s*_4_ and branch rate *r*_4_, where *s*_4_ is the overall evolutionary rate of the subtree originating at node 4 (contains taxon 1 and 2) and *r*_4_ is the evolutionary rate on branch *b*_4_ (**Fig. 7a**). Without assuming a specific distribution of rates, *s*_4_/*r*_4_ cannot be determined uniquely and *t*_4_ can be at any point between 0 and *t*_5_. **Figure 7c** and **7d** present two extreme possibilities. In one, if the subtree rate (*s*_4_) is much higher after the divergence event at node 4 (*s*_4_ >> *r*_4_), then the estimate of *t*_4_ will be small and the divergence event recent (**Fig. 7c**). Alternately, if the subtree rate is much slower after the divergence event at node 4 (*s*_4_ << *r*_4_), then *t*_4_ will be much more ancient (**Fig. 7d**).

In its mathematical formulation, RRF considers the best estimate of the rate of evolution of an ancestral lineage to be the average of the rate of evolution of its two descendant lineages (e.g., equations [3], [14], [15], and [16]), as well as the relative rates among lineages. In the current example, this would result in the timetree shown in **Fig. 7b**. This is the principle of minimum rate change from the ancestor to its immediate descendants, where we do not favor extreme rate assignments, e.g., those in **Fig. 7c** and **Fig. 7d**. Probabilities of such extreme rate assignments are also low in commonly used branch rate distributions (e.g., lognormal, normal, and exponential distributions), so Bayesian methods also tend to favor the smallest rate change needed to explain the data.

### Relationship of RRF with other molecular clock method

The treatment of evolutionary rates is conceptually different between Bayesian methods and RRF, because RRF does not assume a specific statistical distribution for modelling (lineage) rate variation at the outset and Bayesian methods model branch rates rather than the lineage rates. The application of the principle of minimum rate change in RRF is different from non-parametric and semi-parametric approaches based on the idea of Sanderson (1997), because RRF minimizes *lineage rate* changes rather than the *branch rate* changes from an ancestor to its immediate descendants. Also, RRF does not attempt to estimate a universal penalty for the speed of rate change throughout the tree.

RRF is distinct from strict clock approaches because strict clock methods only apply rate averaging (e.g., equations [3] and [4]) at each node in the tree, but RRF imposes an additional constraint that the ratio of sister lineage rates be the ratio of their lineage lengths (e.g., equations [1] and [2]). These additional constraints relax the strict clock and allow rates to vary throughout the tree. For this reason, RelTime is also different from some molecular clock methods that assume the ratio of ages between two nodes to be proportional to the ratio of their average node-to-tip distances, e.g., Purvis (1995), Britton et al. (2002) and Britton et al. (2007). This assumption is tantamount to assuming equality of rates among lineages, and, thus, a strict molecular clock. For this reason, these approaches require pruning of taxa or lineages for which the rate equality does not apply, e.g., Takezaki et al. (1995). RRF does not assume equality among any lineage rates at any time, and it does not require the removal of rate heterogeneous lineages.

### Statistical distribution of relative lineage rates

Relative lineages rates produced by RRF will show extensive correlation, because the evolutionary rate of a lineage is a function of evolutionary rates of all its descendant lineages. This would result in both local and global correlation, which is expected to be present in phylogenies with autocorrelated as well as independent branch rates. As expected, the analysis of 100 sequence datasets showed correlation between ancestral and descendant lineage rates when branch rates were autocorrelated (median correlation = 0.88) or independent (median correlation = 0.77). Therefore, RRF is fundamentally different from the autocorrelation (branch) rate model of Thorne et al. (1998) as well as the independent (branch) rate model of Drummond et al. (2006), as they deal with branches rather than lineages, as defined here. For this reason, the lineage rates produced by RRF are not directly comparable with those observed for branch rates produced by Bayesian methods.

Even though RRF does not require a statistical distribution of lineage rates at the outset, the resulting estimates of lineage rates may follow a statistical distribution. We examined this relationship in an analysis of 100 ingroup sequence datasets in which branch rates were simulated with lognormal, exponential, or uniform distributions. When branch rates followed a lognormal distribution, the distribution of true lineage rates was also lognormal, as was the distribution of RRF lineage rate estimates (**Fig. 8a-d**). When the branch rates followed an exponential distribution, the RRF and true lineage rates showed a similar distribution (**Fig. 8e**). In the case of a uniform distribution of branch rates, the lineage rates showed a normal-like distribution and the RRF rate estimates were lognormally distributed. All of these results suggest that a flexible lognormal distribution will generally fit the distribution of RRF lineage rates. Importantly, time estimates showed a linear relationship with the true times, with slopes close to 1.0 (**Fig. 8g-i**).

**Figure 8.**
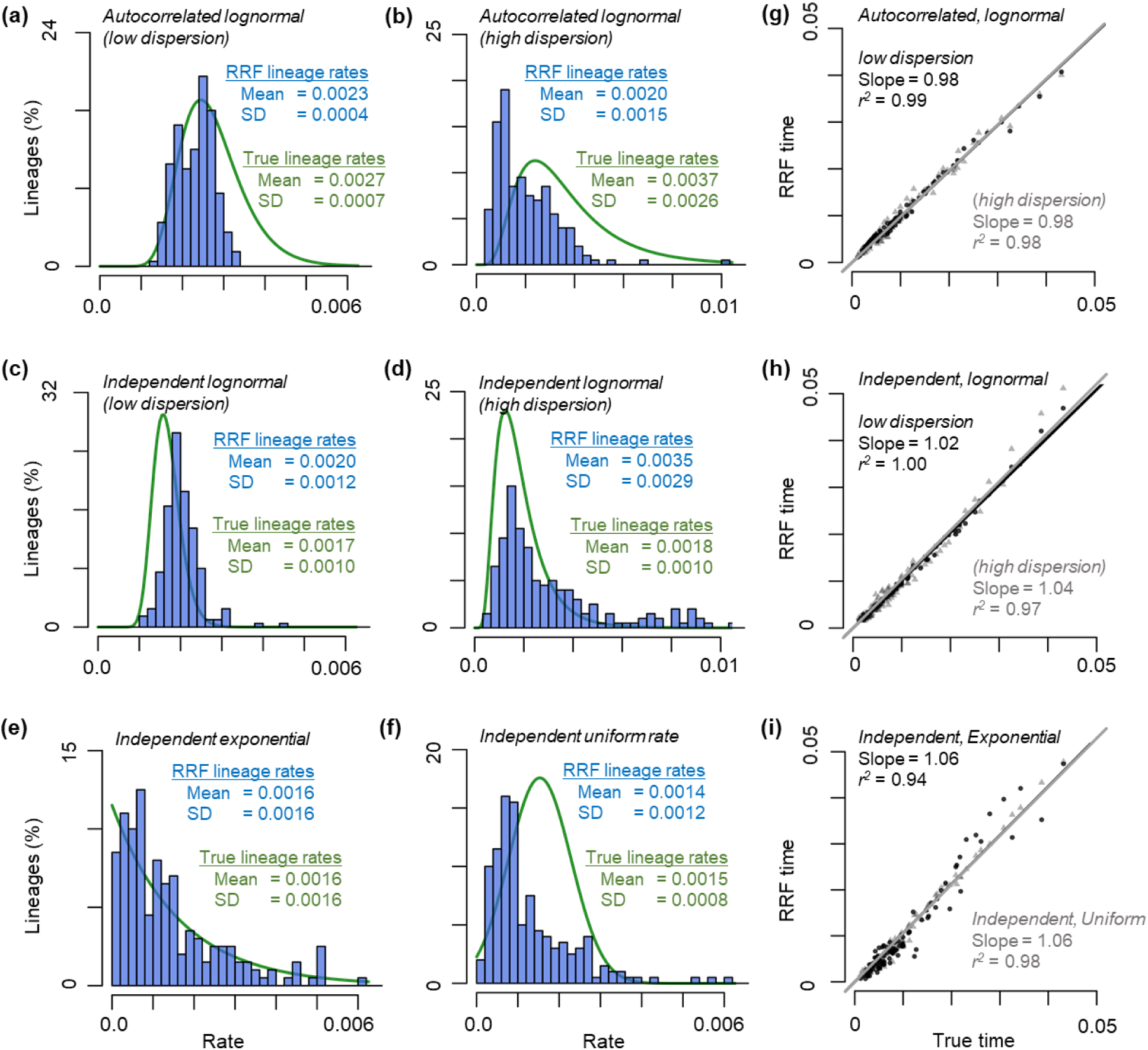
Distributions of true and RRF-derived estimates of lineage rates. Branch rates were simulated under autocorrelated lognormal rate models with (**a**) low and (**b**) high dispersions, independent lognormal rate models with (**c**) low and (**d**) high dispersions, and independent rate models with (**e**) exponential and (**f**) uniform distributions. Green lines represent the fitted curves of the true lineage rate distributions and blue bars show the distributions of RRF rates. Rate unit is substitutions per site per million years. (**g-i**) Relationships between true times and RRF times in six rate scenarios. Black circles and lines represent the average times of 5 replicates simulated under rate scenarios in (**a**), (**c**), and (**e**), and gray triangles and lines represent the average times of 5 replicates simulated under rate scenarios in (**b**), (**d**), and (**f**). All times are normalized to the sum of ingroup divergence times. Regression slopes and correlation coefficients (*r*^2^) are shown.

### Point estimates and their variances

RRF yields point estimates for (relative) lineage rates and divergence times based on branch lengths. As is the common practice in classical statistics, estimates of dispersion such as standard errors and confidence intervals accompany all rate and time estimates. The variance of a lineage rate estimate is a function of branch length variances. It can be obtained analytically by using the equations for lineage rate (e.g., equations [28] - [31] for the case of three taxa with outgroup by the delta method) or simply by using a bootstrap sampling procedure. However, the estimation of confidence intervals around the node ages depends on branch length variances as well as the degree of inequality of evolutionary rates among lineages (Kumar and Hedges 2016). Tamura et al. (2013) proposed a method to estimate confidence intervals within RelTime, which produces rather wide confidence intervals. We are currently investigating an advanced approach to narrow confidence intervals while maintaining appropriate coverage probabilities, but this subject is beyond the scope of the current manuscript.

### Computational speed of RelTime

RRF scales well with increasing numbers of sequences and is much faster than Bayesian methods for analyses of molecular sequence data (**Fig. 1**). The fast computational speed is due to the innovation that RRF uses all the data first to map a large alignment onto a phylogeny, and then it uses the resulting branch lengths to generate relative divergence times and evolutionary lineage rates. The computational time taken by RRF is the sum of time taken to generate maximum likelihood estimates of branch lengths for a given sequence alignment and phylogeny and the time taken to estimate rates and dates using RRF. The latter is negligible compared to the former, because of the analytical nature of RRF. In comparison, Bayesian methods are computationally demanding because they require a substantial exploration of likelihood space using prior distributions to generate posterior estimates of rates and divergence times.

### RelTime for phylogenies with branch lengths

The above decomposition has a positive side effect, in addition to making RelTime computationally speedy. RRF is applicable for any phylogeny where branch lengths reflect the amount of change. For example, RRF is directly applicable when branch lengths are estimated by using pairwise evolutionary distances and a least squares approach for a given tree topology (Rzhetsky and Nei 1993). In addition to multiple sequence alignments, such distances can come from unaligned and locally aligned genome or genomic segments (e.g., (Otu and Sayood 2003; Henz et al. 2004; Auch et al. 2006; Deng et al. 2006; Gao and Qi 2007; Lin et al. 2009; Xu and Hao 2009).

In fact, RRF can be applied to any phylogeny where branch lengths are generated using other types of molecular data (e.g., gene expression patterns and breakpoint distances) or non-molecular data (e.g., morphological and life history traits), e.g., Herniou et al. (2001), Gramm and Niedermeier (2002), King et al. (2016), and Cooney et al. (2017). Of course, the accuracy of the relative rate and time inferences made for such data depends directly on the accuracy of the phylogenetic tree and the branch lengths, so the biological interpretations of the results obtained will require utmost care.

### Usefulness of relative node ages

RRF’s accurate estimation of relative node ages without assuming a speciation-model or calibration priors can benefit many applications (Tamura et al. 2012). For example, relative node ages from molecular data are directly comparable with those from fossil data. This allows evaluation of biological hypotheses without the circularity created by the use of calibration priors and densities inferred from molecular data (Battistuzzi et al. 2015; Gold et al. 2017). Along these lines, RRF has been used to develop a protocol to identify calibration priors that have the strongest influence on the final time estimates in Bayesian dating (Battistuzzi et al. 2015), because the cross-validation methods are unlikely to be effective (Warnock et al. 2012; Warnock et al. 2015).

### Inferring absolute times from relative node ages

By placing calibration constraints on one or more nodes in the tree, we can generate an absolute timetree from the ultrametric tree containing relative node ages. Tamura et al. (2013) presented an algorithmic approach for adjusting relative rates to ensure that the estimated times for calibrated nodes are within researcher-specified boundaries. This process respects maximum and minimum boundaries only, which is preferable when the uncertainty distribution of calibrations are not known precisely. Otherwise, there is high probability of biased time estimation (Hedges and Kumar 2004; Ho and Phillips 2009; Inoue et al. 2010; Heath et al. 2014; Ho and Duchêne 2014; dos Reis et al. 2015). Tamura et al.’s approach worked well in the analysis of large datasets, because Bayesian time estimates reported in multiple large-scale studies are similar to those produced by RRF using ultrametric trees with relative times that were transformed into timetrees using many calibration constraints (Mello et al. 2017).

## Conclusions

We have presented a mathematical foundation for the RelTime method and elucidated its relationship with other relaxed and strict clock methods. We have shown that the relative rate framework (RRF) produces excellent estimates of rates and divergence times for evolutionary lineages. It is, however, important to note that estimates of divergence times in a phylogeny are only biologically meaningful when reconstruction of evolutionary relationships is robust. Therefore, the best practice is to first obtain a reliable phylogeny and then estimate divergence times. We must also consider the confidence intervals associated with node ages to assess the precision of time estimates prior to making biological inferences.

## MATERIALS AND METHODS

### Computer simulations and analysis

We simulated 200 multisequence alignments: 50 each for two models of evolutionary rates (independent and autocorrelated among branches) for two model topologies containing three- and four-ingroup taxa (**Fig. 2a** and **2b**, respectively). The node height of the ingroup subtree was set to be 10 time units, while the node heights of all descendent subtrees varied randomly between 0 and 10 time units. For each resulting model timetree, branch rates were sampled from a lognormal distribution, where the mean rate was drawn randomly from an empirical distribution (Rosenberg and Kumar 2003) and the standard deviation varied from 0.25 to 0.75 for all branches independently. For the autocorrelated rates, the initial rate was drawn randomly from an empirical distribution (Rosenberg and Kumar 2003) and the autocorrelation parameter was varied from 0.1 to 0.3. This rate sampling resulted in a phylogram with branch lengths that could be used as input for SeqGen (Grassly et al. 1997). We used the Hasegawa-Kinshino-Yano (HKY) model (Hasegawa et al. 1985) with 4 gamma categories and empirically-derived GC content and transition/transversion ratio (Rosenberg and Kumar 2003) as simulation parameters. We generated simulated multispecies alignments where each sequence was 3,000 base pairs long. The results from three-ingroup sequence analysis (**Fig. 4**) were similar to those from the analysis of four-ingroup sequences (not shown).

Using the same simulation strategy, we created 35 alignments each under independent and autocorrelated rate scenarios following a master phylogeny of 100 taxa that was sampled from the bony-vertebrate clade in the Timetree of Life (Hedges and Kumar 2009). In the independent rate case, the standard deviation varied from 0.3 to 0.5. In the autocorrelated rate case, the autocorrelation parameter varied from 0.01 to 0.04. All other simulation parameters (GC contents, transition/transversion ratio and sequence length) were derived from empirical distributions (Rosenberg and Kumar 2003).

Using the same 100 ingroup taxa master phylogeny of bony vertebrates, we simulated five additional sequence datasets under (1) an autocorrelated lognormal rate model with low dispersion, where the initial rate and the autocorrelation parameter were set to be *r* and 0.02, respectively; (2) an autocorrelated lognormal rate model with high dispersion, where the initial rate and the autocorrelation parameter were set to be *r* and 0.05, respectively; (3) an independent lognormal rate model with low dispersion, where the mean rate and the standard deviation (in log scale) were set to be *r* and 0.25, respectively; (4) an independent lognormal rate model with high dispersion, where the mean rate and the standard deviation (in log scale) were set to be *r* and 0.75, respectively; (5) an independent exponential rate model, where the mean rate was set to be *r*; and (6) an independent uniform rate model, where the branch-specific rates were sampled from an uniform distribution from 0 to 2*r*. GC contents, transition/ transversion ratio, sequence length and evolutionary rate *r* were derived from empirical distributions (Rosenberg and Kumar 2003).

MEGA software (Kumar et al. 2012; Kumar et al. 2016) was used to obtain maximum likelihood estimates of branch lengths from simulated sequence alignments, where the correct substitution model and tree topology were used. RRF (equations [28] - [42]) was applied to the resulting phylogram with branch lengths, and relative lineage rates and times were obtained. One calibration (true age ± 10Ma) at the crown node of the ingroup was used to convert relative time estimates for comparison with true times (**Fig. 8**). No calibrations were used in other RRF analyses.

All Bayesian analyses were conducted using MCMCTree (Yang 2007) using correct priors; two independent runs of 5,000,000 generations were carried out. Results were checked for convergence using Tracer (Rambaut et al. 2014). ESS values were higher than 200 after removing 10% burn-in samples in each run. MCMCTree analyses used one root calibration (true age ± 0.1 time units).

### Analysis of hybrid rate models

Simulated datasets and BEAST results were provided by Beaulieu et al. (Beaulieu et al. 2015), or retrieved from the Dryad Repository. All outgroup and root calibrations are automatically disregarded in RelTime, because the assumption of equal rates of evolution between the ingroup and outgroup sequences is not testable (Kumar et al. 2016). Lognormal distributions with fixed median values of “true ages” were used as calibration densities in the original study (Beaulieu et al. 2015). Because RelTime does not require specific density distributions for calibrations, we used true age ± 5Ma for all 15 ingroup calibrated nodes in the re-analysis to directly compare RelTime divergence time estimates with those from BEAST. Calibrations employed in RelTime (true age ± 5Ma) had boundaries similar to 99% probability densities of lognormal distributions originally employed as calibrations. The same alignments, topology and ingroup calibrations were used in RRF analyses. Estimates of angiosperm age were obtained by summarizing estimates of 100 datasets each in which the herbaceous clades have 3 times (3x) and 6 times (6x) higher rates than those of woody clades.

## ACKNOWLEDGEMENTS

We are thankful to Dr. Heather Rowe for editorial comments and Dr. Beatriz Mello for many helpful discussions. We thank Jeremy Beaulieu for providing simulated data and Bayesian results. This research was supported by grants from NIH (R01GM126567-01), National Aeronautics and Space Administration (NASA, NNX16AJ30G), and Tokyo Metropolitan University (DB105).

